# Investigating the influence of extraction methodology, and certain other experimental parameters on bacterial response to selected quorum-modulatory plant extracts

**DOI:** 10.1101/511733

**Authors:** Pooja Patel, Chinmayi Joshi, Vijay Kothari

## Abstract

In context of the global threat of antimicrobial resistance (AMR), a large number of labs throughout the world are investigating plant extracts for their possible anti-pathogenic efficacy. Outcome of protocols employed for screening the plant extracts for such activities are influenced by multiple factors. Among them, this study has found the method used for extraction, size and volume of the assay system, to be important factors. Further we demonstrate the extracts prepared by microwave-and vacuum-assisted extraction methods to be stable with respect to their anti-pathogenic efficacy over a long period of storage.

## I. Introduction

Medicinal plants are currently being actively investigated as a potential source of therapeutic phytochemicals that may lead to the development of novel drugs, cosmetics or nutraceuticals [1]. Biological activities of plant-based formulations are significantly dependent on extraction solvent and technique used. With a notable variety of methods present, selection of proper extraction method needs meticulous evaluation [1]. Further, same extract when tested for a particular biological activity using different protocols, results may vary. Present study attempted to investigate effect of certain important experimental factors on outcome of experiments aimed at assessing anti-virulence potential of plant extracts.

## II. Materials and Methods

### A. Plant materials

Peel of *Punica granatum* L. and Seeds of *Phyllanthus emblica* were procured (during 2015-16) from the fruits purchased from local market in the city of Ahmedabad and stored in air tight jar. They were authenticated for their unambiguous identity by Dr. Archana Mankad, Botany Dept., Gujarat University, Ahmedabad. Panchvalkal formulation (PentaphyteP-5^®^) was procured from Dr. Palep’s Medical Research Foundation Pvt. Ltd., Mumbai.

### B. Test organisms

*Chromobacterium violaceum* (MTCC 2656), were procured from MTCC (Microbial Type Culture Collection, Chandigarh). *Pseudomonas aeruginosa* was sourced from our internal culture collection. Nutrient broth and Pseudomonas broth (HiMedia, Mumbai) were used as growth media for *C. violaceum* and *P. aeruginosa* respectively. Incubation temperature and time for both these bacteria were kept at 37 °C for 22-24 h.

### C. Extraction

Dried peels and seeds were powdered to a uniform particle size. Sample to solvent ratio was kept constant for all the methods- 1 g seed powder in 50 ml solvent. 50% ethanol (Ureca consumers co. op stores Ltd, Ahmedabad) was used as extraction solvent.

### D. Microwave assisted extraction (MAE)

It was carried out in a microwave oven (Electrolux EM30EC90SS) at 720 W with intermittent cooling (each cooling cycle was of 40 s) in 250 mL screw capped glass bottle (Borosil). Dark (brown) bottles were used to limit effect of light on plant material. Cap of the bottle was kept little loose during extraction. Total duration of microwave heating for extraction in ethanol was 70s.

### E. Vacuum assisted extraction (VAE)

It was carried out at a working pressure of 7.36 psi. Total duration of extraction time was 15 m in at 65 °C.

Extracts were clarified by centrifugation (Remi BZCI-8729) at 10,000 rpm for 15 min, followed by filtration with Whatman # 1 filter paper (Whatman International Ltd., England). After evaporation, dried extracts were reconstituted in DMSO (Merck, Mumbai). Extraction efficiency (Table 1) was calculated as percentage weight of the starting dried plant material. Reconstituted extracts were stored in autoclaved glass vials (15 mL, Merck) under refrigeration (4-8°C). Internal surface of the vial cap was covered with aluminium foil to prevent the leaching of any compounds from the cap into the extract [2].

**Table 1.** Extraction efficiency of extracts prepared by both methods

|  | *P. emblica* |  | *P. granatum* |  |
| --- | --- | --- | --- | --- |
| Method used for extraction | VAE | MAE | VAE | MAE |
| Extraction efficiency (%) | 10.30 | 12.40 | 35.30 | 28.20 |
| Reconstitution efficiency (%) | 84.46 | 88.71 | 93.48 | 94.32 |
| Final Concentration after reconstitution (mg/mL) | 56.86 | 71.10 | 221.62 | 172.05 |

### F. Broth dilution assay

Effect of test extracts on bacterial growth and production of quorum sensing (QS)-regulated pigments were assessed using the methods described by us previously [3]. Appropriate vehicle control containing DMSO was also included in the experiment, along with abiotic control (containing extract, but no inoculum). Catechin was used as positive control.

### G. In vivo assay

*In vivo* efficacy of the test extracts was evaluated using the nematode worm *Caenorhabditis elegans* as the model host through live-dead assay as detailed by us in Patel et al. [4]. Standard antibiotics-and catechin-treated bacterial suspensions were used as positive control.

### H. Statistical analysis

All the experiments were performed in triplicate, and measurements are reported as mean ± standard deviation (SD). Statistical significance of the data was evaluated by applying *t*-test using Microsoft Excel^®^. *p* values less than 0.05 were considered to be statistically significant.

## III. RESULTS

### A. Choice of extraction method can have significant influence on anti-infective activity

*P. granatum* and *P. emblica* extracts prepared by MAE and VAE methods were assessed for their ability of influence bacterial growth and QS-regulated pigment production *in vitro*, and bacterial virulence towards *C. elegans in vivo*. Against *P. aeruginosa* both plant extracts prepared by any of the method had identical *in vitro* effect. However, in case of *P. emblica*, extract prepared by VAE could attenuate virulence of *P. aeruginosa* more than that prepared by MAE; whereas in case of *P. granatum*, extract prepared by MAE method scored better during *in vivo* assay (Figure 1). Against *C. violaceum*, though the extracts prepared by both methods differed with respect to their in vitro effect, their *in vivo* efficacies were equivalent (Figure 2).

**Figure 1:**
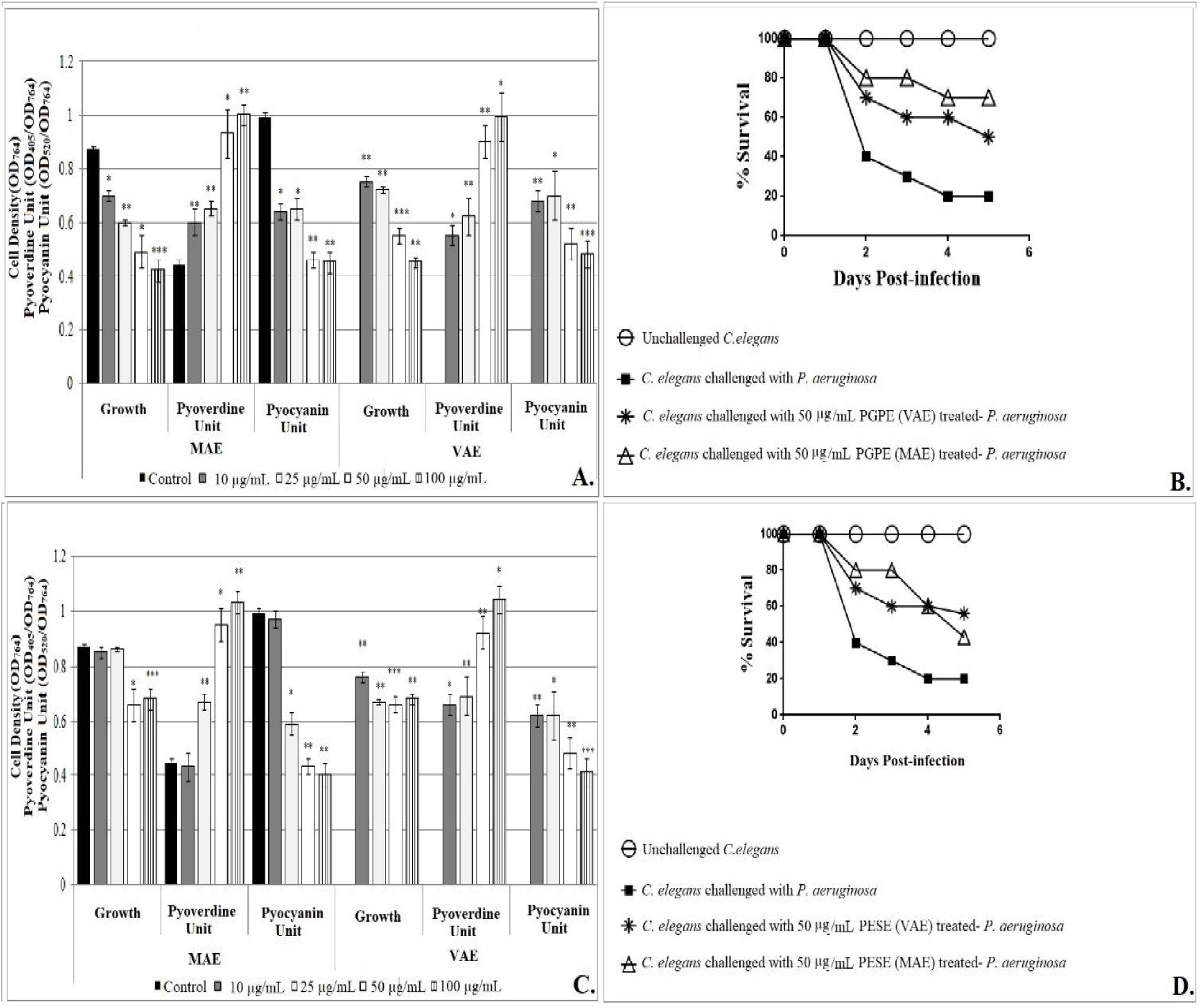
Comparison of *in vitro* and *in vivo* efficacy of plant extracts prepared by VAE and MAE against *P. aeruginosa*. (A) Effect of PGPE on growth and QS-regulated pigment production in *P. aeruginosa* (B) PGPE prepared by MAE scored better *in vivo* anti-virulence effect than that prepared by VAE (C) Effect of PESE on growth and QS-regulated pigment production in *P. aeruginosa* (D) PESE prepared through both extraction methods were statistically at par with respect to their *in vivo* efficacy During *in vitro* experiments, bacterial growth was measured as OD_764_; OD of pyoverdine was measured at 405 nm, Pyocyanin was measured at 520 nm. Pyoverdine Unit was calculated as the ratio OD_405_/OD_764_ (an indication of pyoverdine production per unit of growth); Pyocyanin Unit was calculated as the ratio OD_520_/OD_764_ (an indication of pyocyanin production per unit of growth); Catechin (50 µg/mL) inhibited pyoverdine 17.13%** ± 0.06 and pyocyanin 23.65%* ± 0.04 production without affecting the bacterial growth. During *in vivo* experiments, DMSO present in the ‘vehicle control’ at 0.5%v/v did not affect virulence of the bacterium towards *C. elegans*. DMSO (0.5%v/v) and plant extracts at tested concentrations showed no toxicity towards the worm.

**Figure 2:**
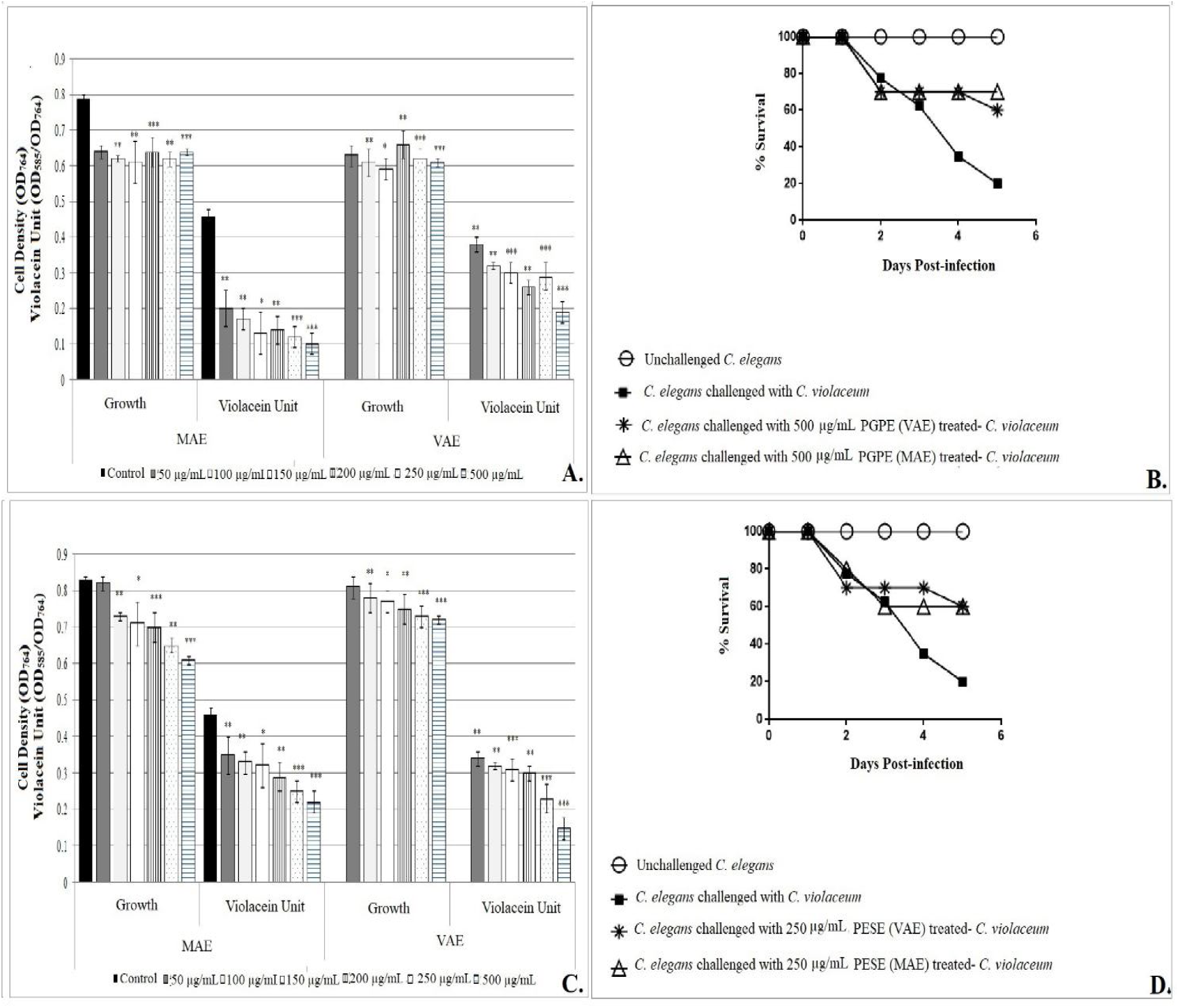
Comparison of *in vitro* and *in vivo* efficacy of plant extracts prepared by VAE and MAE against *C. violaceum*. (A) Effect of PGPE on growth and QS-regulated pigment production in *C. violaceum* (B) PGPE prepared through both extraction methods were statistically at par with respect to their *in vivo* efficacy (C) Effect of PESE on growth and QS-regulated pigment production in *C. violaceum* (D) PESE prepared through both extraction methods were statistically at par with respect to their *in vivo* efficacy During *in vitro* experiments, Bacterial growth was measured asOD_764_; OD of violacein was measured at 585 nm, and Violacein Unit was calculated as the ratio OD_585_/OD_764_ (an indication of violacein production per unit of growth); Catechin (50 µg/mL) did not exerted any effect on growth of *C. violaceum*, and inhibited violacein production by 47.69%*** ± 0.03. During *in vivo* experiments, DMSO present in the ‘vehicle control’ at 0.5%v/v did not affect virulence of the bacterium towards *C. elegans*. DMSO (0.5%v/v) and plant extracts at tested concentrations showed no toxicity towards the worm. Catalase assay was done by monitoring disappearance of H_2_O_2_ at 240 nm; Hemoglobin concentration was measured at OD_540._

### B. Vessel size and reaction volume are important determinants of experimental output

Effect of *Panchvalkal* formulation (PF) on *C. violaceum* and *P. aeruginosa* (Figure 3) was assessed in two different sized vessels: 10 mL test tubes (reaction volume 1 mL), and 250 mL flask (reaction volume 100 mL). PF could influence bacterial growth, catalase and haemolytic activity, and pigment production to a greater extent when experiments were performed in larger vessel size with higher reaction volume. Similar observation could be made while assessing effect of PF on *P. aeruginosa* catalase and haemolytic activity; however pyocyanin production was affected by PF more in tubes than in flasks.

**Figure 3:**
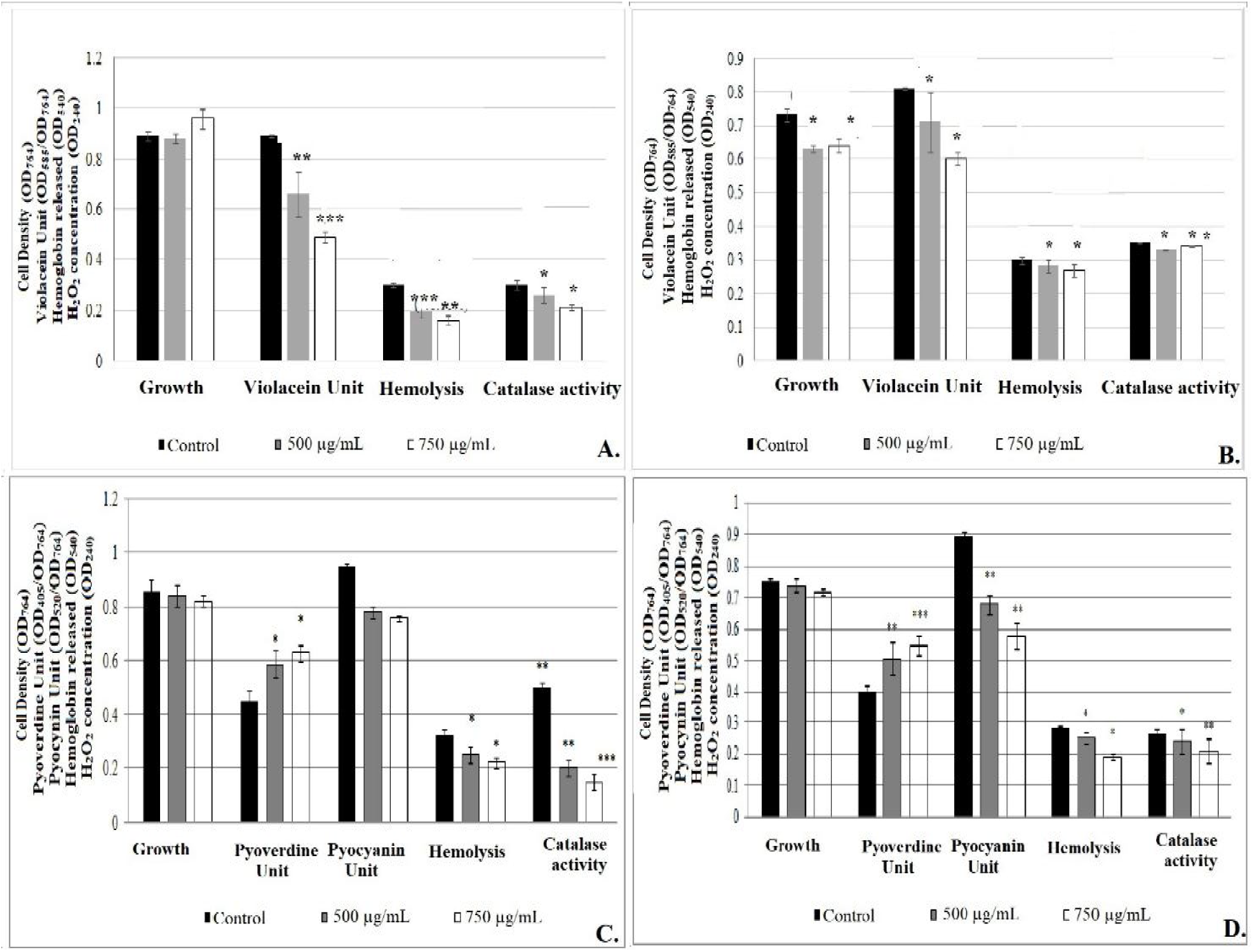
Influence of vessel size and reaction volume on PGPE’s effect on various traits of *C. violaceum* and *P. aeruginosa*. Effect of PGPE on *C. violaceum* grown in (A) flask, and (B) test tube Bacterial growth was measured asOD_764_; OD of violacein was measured at 585 nm, and Violacein Unit was calculated as the ratio OD_585_/OD_764_ (an indication of violacein production per unit of growth); Catechin (50 µg/mL) did not exerted any effect on growth of *C. violaceum*, and inhibited violacein Effect of PGPE on *P. aeruginosa* grown in (C) flask and (D) test tube Bacterial growth was measured as OD_764_; OD of pyoverdine was measured at 405 nm, Pyocyanin was measured at 520 nm. Pyoverdine Unit was calculated as the ratio OD_405_/OD_764_ (an indication of pyoverdine production per unit of growth); Pyocyanin Unit was calculated as the ratio OD_520_/OD_764_ (an indication of pyocyanin production per unit of growth); Catechin (50 µg/mL) inhibited pyoverdine 17.13%** ± 0.06 and pyocyanin 23.65%* ± 0.04 production without affecting the bacterial growth.

### C. Extracts prepared using MAE and VAE method retain their efficacy over long term

*P. granatum* peel extract (PGPE) prepared by MAE, and *P. emblica* seed extract (PESE) prepared by VAE were first prepared and tested for their anti-pathogenic activity in May 2015. Then they were kept stored in refrigerator (4-8 °C) and re-tested in Feb 2016 and Jan 2017 to investigate whether they still have retained their efficacy. *In vitro* and *in vivo* experiments (Figure 4) confirmed that these extracts were stable in terms of their activity against *C. violaceum* over a period of almost 19 months.

**Figure 4:**
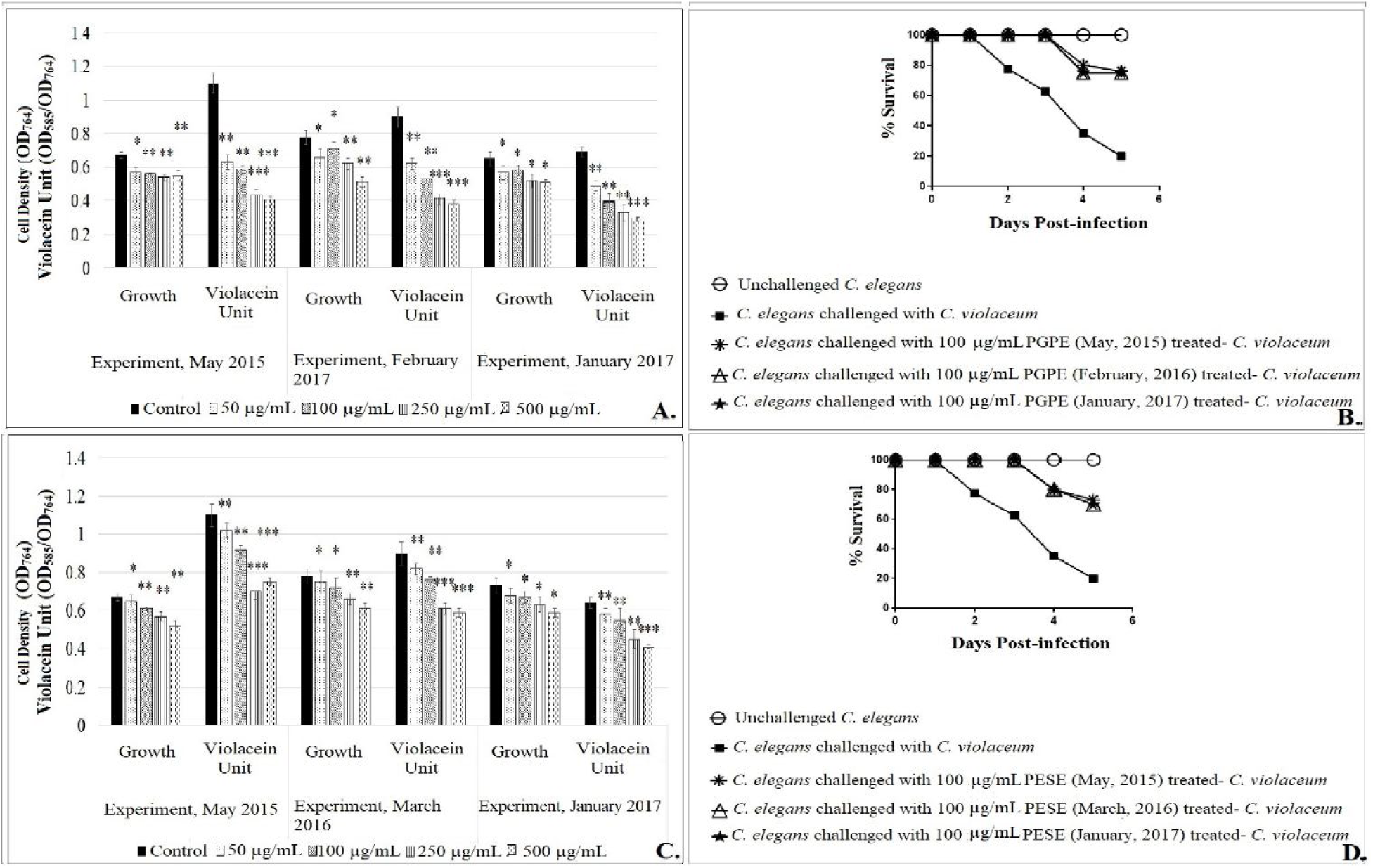
Extracts prepared by VAE and MAE retain their anti-pathogenic effect over a long period of refrigerated storage. A-B. Results for PGPE; C-D. Results for PESE

## IV. Conclusion

Currently plant sources are being actively investigated for a variety of biological and pharmacological activities in search of novel leads of pharmaceutical relevance. However owing to a variety of screening protocols being used for preliminary assessment of desired biological activity, direct comparison between results of different protocols remains tricky. This study has indicated that due attention should be paid to factors like choice of extraction method, vessel size, reaction volume, etc. while fixing the screening strategy. This study has also found the extracts prepared by MAE and VAE method to be stable over a refrigerated storage of over a year. Owing to their inherent character of offering fast heating (MAE), and boiling at lower temp (VAE), these methods are also considered suitable for extraction of heat-labile phytocompounds. Such studies with more varieties of extraction methods, on other experimental determinants, and additional pathogenic bacteria will provide better insight enabling the researchers to adopt potentially more reliable and reproducible screening protocols.

## ACKNOWLEDGMENT

Authors thank Nirma Education and Research Foundation (NERF, Ahmedabad) for financial and infrastructural support

